# A BERT-Based Hybrid System for Chemical Identification and Indexing in Full-Text Articles

**DOI:** 10.1101/2021.10.27.466183

**Authors:** Arslan Erdengasileng, Keqiao Li, Qing Han, Shubo Tian, Jian Wang, Ting Hu, Jinfeng Zhang

**Affiliations:** Florida State University: Department of Statistics, Tallahassee, USA; Cloudmedx: Engineer team, Palo Alto, USA

**Keywords:** chemical NER, chemical normalization, chemical indexing, BERT, sieve-based dictionary matching

## Abstract

Identification and indexing of chemical compounds in full-text articles are essential steps in biomedical article categorization, information extraction, and biological text mining. BioCreative Challenge was established to evaluate methods for biological text mining and information extraction. Track 2 of BioCreative VII (summer 2021) consists of two subtasks: chemical identification and chemical indexing in full-text PubMed articles. The chemical identification subtask also includes two parts: chemical named entity recognition (NER) and chemical normalization. In this paper, we present our work on developing a hybrid pipeline for chemical named entity recognition, chemical normalization, and chemical indexing in full-text PubMed articles. Specifically, we applied BERT-based methods for chemical NER and chemical indexing, and a sieve-based dictionary matching method for chemical normalization. For subtask 1, we used PubMedBERT with data augmentation on the chemical NER task. Several chemical-MeSH dictionaries including MeSH.XML, SUPP.XML, MRCONSO.RFF, and PubTator chemical annotations are used in a specific order to get the best performance on chemical normalization. We achieved an F1 score of 0.86 and 0.7668 on chemical NER and chemical normalization, respectively. For subtask 2, we formulated it as a binary prediction problem for each individual chemical compound name. We then used a BERT-based model with engineered features and achieved a strict F1 score of 0.4825 on the test set, which is substantially higher than the median F1 score (0.3971) of all the submissions.

## I. Introduction

Named entity recognition (NER) is a crucial task in biological natural language processing (NLP) that extracts relevant names from a given corpus. It is usually the first step in many NLP pipelines. Among all the entity types, chemical entities are one of the most searched terms in the PubMed database (1). There have been many methods developed in the past for chemical NER and deep learning methods such as BiLSTM (2), Spacy (3), and OSCAR4 (4) have substantially improved the performance compared to traditional methods. Recently, BERT and its variants (5–10) have achieved the state-of-the-art (SOTA) performance.

BioCreative Challenge was established in 2004 to evaluate information extraction (IE) and text mining (TM) systems applied to the biological domain (11). In each round (every two years), several tracks are organized for different information extraction or text mining tasks. In BioCreative IV and V chemical NER tracks were organized where the corpora contain only the titles and abstracts of PubMed articles. As more full-text articles become publicly available at PubMed Central (PMC), NER methods for full-text articles are needed for biological NLP tasks to take advantage of this resource. NER for full-text presents some different challenges: (1) while titles and abstracts contain only important chemical names in the articles, full-texts contain a more diverse set of chemical names, often with different contexts (i.e., chemical names mentioned in experimental protocols); (2) full-texts contain more abbreviations than titles and abstracts, whose full names are only mentioned in the early part of the articles. Abbreviations can represent totally different concepts in different contexts; and (3) full-texts often contain more noise, such as grammatical errors and typos. In addition, the writing style for full-texts is very different from that of the titles and also different from abstracts since there is often a stricter limit on the number of words on abstracts. In track 2 of BioCreative VII held in the summer of 2021, full-text articles annotated by experts (1) were provided to the participants to evaluate chemical NER methods developed for full-texts. The corpus is called the NLM-Chem dataset, which includes 150 PubMed articles with annotated chemical mentions, corresponding MeSH terms, and the MeSH terms used for indexing the articles.

In BioCreative VII challenge, we first experimented with several BERT-based models, including BERT large uncased (5), BioBERT (7), BlueBERT (8), SciBERT (6), ClinicalBERT (9), and PubMedBERT (10). PubMedBERT outperformed other models when evaluated using the development data. BioBERT was used by the organizer as the baseline model with an F1 score of 0.803. We also tried several data augmentation methods, which helped to further improve the F1 score of the NER task to 0.86.

Chemical normalization is a necessary step after NER, where the goal is to link the identified chemical mentions to the correct MeSH terms. Synonyms of the same entity will all be linked to an official name. In BioCreative VII, chemical normalization is the second part of the chemical identification subtask. For chemical normalization, we used several dictionaries, such as MeSH, supplementary concept records, UMLS MRCONSO, PubTator annotations, and NLM-Chem golden standard dataset, in a specific order and achieved an F1 score of 0.7668 on the test set.

Chemical indexing subtask in BioCreative VII Track 2 is a task that predicting which identified MeSH terms in the full-text PubMed articles should be indexed. We participated in the subtask unofficially (submitted our blind-test result after the deadline). We tried two different approaches. The first approach formulates the problem as an extreme multi-label classification problem to predict all indexed MeSH terms simultaneously using full-texts as input. The second approach formulates the problem as a binary prediction by predicting each MeSH at a time using information extracted for that MeSH term together with engineered features. We used PubMedBERT for the second approach.

In our experiment, the first approach achieved an F1 score of 42% on the development set, and the second approach achieved an F1 score of 61.9% on the development set.

## II. Method

BioCreative VII Track 2 consists of two subtasks, chemical identification, and chemical indexing. The first subtask includes two problems, chemical named entity recognition (NER) and chemical normalization. We developed 3 different systems for each task.

### A. Chemical Named Entity Recognition

As BERT-based models achieved SOTA performance on NER tasks, we first tried several variants of the BERT model including BioBERT, PubMedBERT, SciBERT, BlueBERT, ClinicalBERT, and BERT-large-uncased.

The performance of these models on the test set is shown in Table I. Among the tested models, PubMedBERT achieved the highest F1-score on the development set and test set. PubMedBERT is a BERT model that pre-trained on biomedical text from the scratch by Microsoft research team. The assumption is that pre-training the BERT model solely on the domain-specific text would perform better than general-domain text (10). PubMedBERT outperformed all prior language models and obtained new SOTA results in a wide range of biomedical applications (10). We chose to use PubMedBERT as the base model for chemical NER task.

**TABLE I.** BERT-based Model Performance

| Model | Val F1 | Test F1 | Strict-P | Strict-R | Strict-F |
| --- | --- | --- | --- | --- | --- |
| BioBERT | 0.8462 | 0.8433 | 0.7853 | 0.8068 | 0.7959 |
| BlueBERT | 0.8455 | 0.8442 | 0.776 | 0.826 | 0.8002 |
| SciBERT | 0.8621 | 0.8495 | 0.7855 | 0.8245 | 0.8045 |
| PubMedBERT | 0.8773 | 0.8679 | 0.8085 | 0.8419 | 0.8249 |
| ClinicalBERT | 0.7818 | 0.8114 | 0.7679 | 0.7666 | 0.7672 |
| BERT-large-uncased | 0.8233 | 0.8018 | 0.7732 | 0.7514 | 0.7622 |

BioCreative VII also provided CHEMDNER and BC5CDR datasets from BioCreative IV and V for chemical identification tasks. We trained models on different combinations of the datasets to see if adding more data would increase the performance. Three following models were trained on NLM-Chem training set only, NLM-Chem training set + BC5CDR set, and NLM-Chem training set + BC5CDR + CHEMDNER.

The validation set and test set for the three different models were the same: NLM-Chem validation set and NLM-Chem test set. The results were shown in Table II.

**TABLE II.** Performance with Different Datasets

| Model | Val F1 | Test F1 | Strict-F | Dataset |
| --- | --- | --- | --- | --- |
| PubMedBERT | 0.8773 | 0.8679 | 0.8249 | NLM |
| PubMedBERT | 0.8841 | 0.8617 | 0.817 | NLM+CDR, NLM |
| PubMedBERT | 0.8676 | 0.8115 | 0.7686 | NLM+CDR+CHEMDNER, NLM |

**TABLE III.** Chemical Mention Recognition

| File | Strict-P | Strict-R | Strict-F |
| --- | --- | --- | --- |
| Run 1 | 0.8544 | 0.8658 | 0.8600 |
| Run 2 | 0.8643 | 0.8403 | 0.8521 |
| Run 3 | 0.8440 | 0.7896 | 0.8159 |
| Run 4 | 0.8457 | 0.8617 | 0.8536 |

**TABLE IV.**
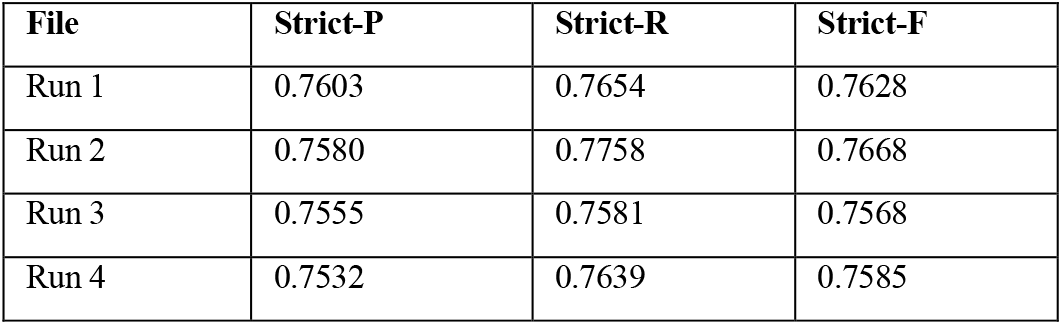
Chemical Normalization to Mesh

| File | Strict-P | Strict-R | Strict-F |
| --- | --- | --- | --- |
| Run 1 | 0.7603 | 0.7654 | 0.7628 |
| Run 2 | 0.7580 | 0.7758 | 0.7668 |
| Run 3 | 0.7555 | 0.7581 | 0.7568 |
| Run 4 | 0.7532 | 0.7639 | 0.7585 |

This experiment showed that adding more data from BioCreative IV and V did not improve the NER performance. One possible reason is that CHEMDNER and BC5CDR annotations may have systematic differences from NLM-Chem for two reasons: CHEMDNER and BC5CDR were extracted from abstracts only, while NLM-Chem only annotates the chemical mentions in full-texts; and the annotation rules may also differ for these corpora. Since the final test set is annotated by the same annotators using the same annotation guideline as the NLM-Chem training and validation dataset, we only used the NLM-Chem dataset to train the model.

We then used a data augmentation technique to further improve the model’s performance. Specifically:

1. We replaced the chemical entities with random strings (i.e., Aspirin→badjaxfjfg).
2. We randomly selected one non-chemical entity in sentences which contain chemical entities, then replaced it with a random string (i.e., that →hsw).

We found that this data augmentation procedure improved the performance slightly.

From the PubMedBERT output, we added some postprocessing steps. First, Ab3P (12), an abbreviation definition detector trained on PubMed abstracts, was used to recognize abbreviations in the text. The full names and their abbreviations are linked within the same articles and all the occurrences received the same NER label.

Secondly, chemical names can be part of other entity names. In such cases, the annotation rule does not label such names as chemicals. For example, in “glucose transporter” glucose is a chemical compound, but the whole word is a protein. PubMedBERT sometimes failed in such cases by labeling glucose as a chemical entity. We manually selected some words such as “enzyme”, “transporter” and “receptor” etc., and corrected any wrongly labeled chemical names immediately before these protein names.

Thirdly, which is optional, we trained another BioBERT based protein NER model to detect protein entities. The goal was to further remove wrongly labeled chemical names which are part of protein names. The rule is:

1. If a token word was recognized by PubMedBERT based Chemical NER model as a chemical entity and by BioBERT based protein NER model as a protein entity at the same time, its predicted entity label (“B” or “I”) would be manually changed “O”.
2. If a predicted chemical entity name was followed by a predicted protein entity name, then the predicted chemical entity label (“B” or “I”) was changed to “O”.

### B. Chemical Normalization

Chemical normalization is to link identified chemical mentions to MeSH terms. MeSH stands for Medical Subject Headings and it is a comprehensive controlled vocabulary for biomedical terms, which are used by PubMed for indexing journal articles about life science (13). We built a sieve-based pipeline using multiple dictionaries as follows:

MeSH: MeSH data is an official lexicon that is updated annually. We downloaded the MeSH.XML file from NLM website. We filtered the MeSH data by keeping only chemicals and drugs, whose MeSH IDs start with the letter D.
SCR: Supplementary concept records (SCR) are created for some chemicals, drugs, and other concepts such as rare diseases. SCR begins with the letter C.
MRCONSO: MRCONSO is a metathesaurus that is often used by UMLS, containing terms, term types, and codes. Terms include chemicals and their synonyms, and codes include MeSH terms. The MeSH terms in this file consist of two parts: terms start with the letter D and terms start with the letter C. We call them MRD and MRC in the following text.
PubTator: PubTator is a web-based tool that identifies entities in biomedical articles from PubMed and PMC (14,15). We extracted chemicals and MeSH terms from PubMed articles processed by PubTator.
NLM-Chem: NLM-Chem dataset is the manually curated corpus made for BioCreative Challenge VII.

As shown in Figure 1, there are three steps to map a chemical to a MeSH ID. The first step is to scan through the dictionaries in the following order: PubTator, MRC, SUPP, MRD, MeSH, and NLM, which is arranged by ascending precision. NLM is the most accurate dictionary and PubTator is the least accurate. For terms in more than one dictionary, we replaced the terms mapped earlier with those mapped later; The second step is to use Ab3p to process those unmapped chemicals to detect abbreviations and their long forms; The third step is to scan the detected long forms of abbreviations through the dictionaries in the same order in the first step.

**Fig. 1.**
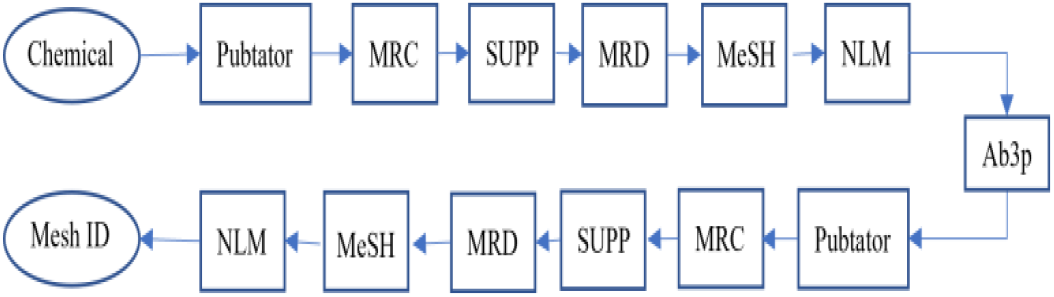
Mesh NormalizatioN FlowChart

### C. Chemical Indexing

Subtask 2 is a chemical indexing prediction task for full-text PMC articles. In this task, a full-text article is given, and a list of chemical MeSH terms should be predicted as the chemical indexing of this article. We tried two different approaches this task.

1. Building an extreme multi-label classification system using a large number of PubMed articles with MeSH indexing. We selected 967,826 PMC full-text articles with chemical indexing information extracted from the PubMed database. For each article, we collected title, abstract, body text (also contains section information), chemical names, and MeSH indexing terms. In our experiment, we fed the model with MeSH terms as labels and title, abstract, and body text as input. Therefore, the MeSH indexing prediction problem became an extreme multi-label classification problem since the total number of possible MeSH indexing terms is very large (16). We tested two models called fastText (15) and extremeText (15,16) to predict the MeSH indexing for the full-text articles. fastText is a library for learning word embeddings and text classification created by Facebook’s AI Research lab (15). It’s dedicated to text classification and learning word representations and was designed to allow for quick model iteration and refinement without specialized hardware. fastText models can be trained on more than a billion words on any multicore CPU in less than a few minutes (15). extremeText is an extension of fastText library for multi-label classification including extreme cases with hundreds of thousands or even millions of labels (16). This model is suitable for our problem because the number of unique labels is around 80,000 and input text data are much larger. It turned out that the best F1 score we could achieve was 0.42, which was not satisfactory for us. Thus, we decided not to submit this result before the official deadline.
2. Building a binary MeSH indexing classification system using a PubMedBERT model with engineered features. In this strategy, we dealt with one MeSH term at a time by predicting whether it should be used for indexing or not. To remove the noise from the long text, we broke up full-texts into sentences and selected the sentences with chemical mentions of the corresponding MeSH terms as input to the model. The labels are simply True or False based on if the MeSH terms were used for indexing the articles or not. We added engineered features before the sentences, such as the section where the sentences were taken from and the chemical names whose MeSH terms were to be predicted. The F1 score of this approach on the NLM-Chem development set was 0.619.

## III. Results

Our result of subtask 1 is quite promising on the final test set. Our best strict F1 score of chemical mention on the test set is 0.86 and the best strict F1 score of chemical normalization is 0.7668. We submitted four runs for chemical mention recognitions and chemical normalization, respectively. Below we provide the details for each run:

Run 1: a. Data augmented by: (1) replacing each of the chemical entities with a random string; (2) selecting 50% of sentences which contain chemical entities and randomly choosing one non-chemical entity and replace it with a random string while the chemical entities remain unchanged; b. Using Ab3P to post-process the prediction results to add chemical entity tags and remove wrong chemical tags; c. Manually selected some “protein” words (i.e., “transporter”) and changed the labels of the corresponding chemical entities, which are immediately before these selected words, to “O”, if they were labeled as “B” or “I”.
Run 2: a. Data augmented by: (1) replacing each of the chemical entities with a random string; (2) selecting 70% of sentences which contain chemical entities and randomly choosing one non-chemical entity and replacing it with a random string while the chemical entities remain unchanged; b. Same as Run 1; c. Same as Run 1.
Run 3: a. Same as Run 2; b. Same as Run 1; c. Same as Run 1; d. Using the BioBERT protein NER model to detect protein entities and changing the label of the chemical entities which are part of a longer protein name to “O” if they were labeled as “B” or “I”.
Run 4: a. Data augmented by: (1) replacing each of the chemical entities with a random string; (2) for all sentences which contain chemical entities and randomly selecting one non-chemical entity and replacing it with random string while the chemical entities remain unchanged; b. Same as Run 1; c. Same as Run 1; d. Same as Run 3.

## IV. CONCLUSION

In this paper, we describe our solutions for three different chemical identification/indexing tasks. We selected PubMedBERT from several BERT-based models and implemented data augmentation technique to further improve the performance. We found out that adding CHEMDNER and BC5CDR corpora to the training data did not improve the NER result on NLM-Chem test set. This was a somewhat surprising result to us. We believe that the community should study their systematic differences so that we will either find out a way to use all of the datasets to train better models, or we can make a recommendation to researchers that certain datasets are problematic and should not be used in the future.

We evaluated several available dictionaries for MeSH normalization and optimized their order for the sequential scanning of the identified terms, which significantly improved the accuracy. The order of the dictionaries maximized the information we can use from different dictionaries. However, as databases and methods are being constantly updated, their quality can also change over time. The optimal order may need to be adjusted accordingly in the future.

Chemical indexing is a challenging task and we tried two different approaches using two different datasets. Our first approach was to use a large amount of PMC PubTator annotations as training set and the second approach was to use NLM-Chem as training set. Even though the first approach has an advantage of using a large dataset, the chemical indexing result was not ideal. The possible reason may be that PubTator annotations are not as accurate as pure human annotations in NLM-Chem data. The different formulations of the problem may have also played a role. Additional modifications (i.e. more engineered features) may be made in the future to the second approach to further improve its performance.

## Acknowledgment

JZ was supported partially by a grant from the National Institute of General Medical Sciences of the National Institutes of Health grant No. R01GM126558.

## References

1. Islamaj R, Leaman R, Kim S, Kwon D, Wei C-H, Comeau DC, et al. NLM-Chem, a new resource for chemical entity recognition in PubMed full text literature. Scientific Data 2021 8:1 [Internet]. 2021 Mar 25 [cited 2021 Oct 5];8(91). Available from: https://www.nature.com/articles/s41597-021-00875-1

2. Huang Z, Xu W, Yu K. Bidirectional LSTM-CRF Models for Sequence Tagging. 2015 Aug 9 [cited 2021 Oct 8]; Available from: https://arxiv.org/abs/1508.01991v1

3. Facts & Figures · spaCy Usage Documentation [Internet]. [cited 2021 Oct 8]. Available from: https://spacy.io/usage/facts-figures

4. Li X, Yin X, Li C, Zhang P, Hu X, Zhang L, et al. Oscar: Object-Semantics Aligned Pre-training for Vision-Language Tasks. Lecture Notes in Computer Science (including subseries Lecture Notes in Artificial Intelligence and Lecture Notes in Bioinformatics) [Internet]. 2020 Apr 13 [cited 2021 Oct 8];12375 LNCS:121–37. Available from: https://arxiv.org/abs/2004.06165v5

5. Devlin J, Chang M-W, Lee K, Toutanova K. BERT: Pre-training of Deep Bidirectional Transformers for Language Understanding. NAACL HLT 2019 – 2019 Conference of the North American Chapter of the Association for Computational Linguistics: Human Language Technologies – Proceedings of the Conference [Internet]. 2018 Oct 11 [cited 2021 Oct 8];1:4171–86. Available from: https://arxiv.org/abs/1810.04805v2

6. Beltagy I, Lo K, Cohan A. SciBERT: A Pretrained Language Model for Scientific Text. EMNLP-IJCNLP 2019 – 2019 Conference on Empirical Methods in Natural Language Processing and 9th International Joint Conference on Natural Language Processing, Proceedings of the Conference [Internet]. 2019 Mar 26 [cited 2021 Oct 8];3615–20. Available from: https://arxiv.org/abs/1903.10676v3

7. Lee J, Yoon W, Kim S, Kim D, Kim S, So CH, et al. BioBERT: a pre-trained biomedical language representation model for biomedical text mining. Bioinformatics [Internet]. 2019 Jan 25 [cited 2021 Oct 8];36(4):1234–40. Available from: https://arxiv.org/abs/1901.08746v4

8. Peng Y, Yan S, Lu Z. Transfer Learning in Biomedical Natural Language Processing: An Evaluation of BERT and ELMo on Ten Benchmarking Datasets. 2019 Jun 13 [cited 2021 Oct 8];58–65. Available from: https://arxiv.org/abs/1906.05474v2

9. Alsentzer E, Murphy JR, Boag W, Weng W-H, Jin D, Naumann T, et al. Publicly Available Clinical BERT Embeddings. 2019 Apr 6 [cited 2021 Oct 8]; Available from: https://arxiv.org/abs/1904.03323v3

10. Gu YU, Tinn R, Cheng H, Lucas M, Usuyama N, Liu X, et al. Domain-Specific Language Model Pretraining for Biomedical Natural Language Processing. 2021 [cited 2021 Oct 5];(1):24. Available from: https://doi.org/10.1145/3458754.

11. L H, A Y, C B, A V. Overview of BioCreAtIvE: critical assessment of information extraction for biology. BMC bioinformatics [Internet]. 2005 May 24 [cited 2021 Oct 8];6 Suppl 1(Suppl 1). Available from: https://pubmed.ncbi.nlm.nih.gov/15960821/

12. Sohn S, Comeau DC, Kim W, Wilbur WJ. Abbreviation definition identification based on automatic precision estimates. BMC Bioinformatics [Internet]. 2008 Sep 25 [cited 2021 Oct 5];9:402. Available from: pmc/articles/PMC2576267/

13. Coletti MH, Bleich HL. Medical Subject Headings Used to Search the Biomedical Literature. Journal of the American Medical Informatics Association : JAMIA [Internet]. 2001 [cited 2021 Oct 9];8(4):317. Available from: pmc/articles/PMC130076/

14. Wei C-H, Kao H-Y, Lu Z. PubTator: a web-based text mining tool for assisting biocuration. Nucleic Acids Research [Internet]. 2013 Jul 1 [cited 2021 Oct 8];41(W1):W518–22. Available from: https://academic.oup.com/nar/article/41/W1/W518/1105731

15. CH W, A A, R L, Z L. PubTator central: automated concept annotation for biomedical full text articles. Nucleic acids research [Internet]. 2019 Jul 1 [cited 2021 Oct 8];47(W1):W587–93. Available from: https://pubmed.ncbi.nlm.nih.gov/31114887/

16. Liu J, Chang W-C, Wu Y, Yang Y. Deep Learning for Extreme Multi-label Text Classification. 2017;

17. Joulin A, Grave E, Bojanowski P, Mikolov T. Bag of Tricks for Efficient Text Classification. 15th Conference of the European Chapter of the Association for Computational Linguistics, EACL 2017 – Proceedings of Conference [Internet]. 2016 Jul 6 [cited 2021 Oct 5];2:427–31. Available from: https://arxiv.org/abs/1607.01759v3

18. Chang W-C, Yu H-F, Zhong K, Yang Y, Dhillon I. Taming Pretrained Transformers for Extreme Multi-label Text Classification. arXiv.org. 2019 May 7;

